# Patterning precision under non-linear morphogen decay and molecular noise

**DOI:** 10.1101/2022.11.04.514993

**Authors:** Jan A. Adelmann, Roman Vetter, Dagmar Iber

## Abstract

Morphogen gradients can instruct cells about their position in a patterned tissue. Non-linear morphogen decay has been suggested to increase gradient precision by reducing the sensitivity to variability in the morphogen source. Here, we use cell-based simulations to quantitatively compare the positional error of gradients for linear and non-linear morphogen decay. While we confirm that non-linear decay reduces the positional error close to the source, the reduction is very small for physiological noise levels. Far from the source, the positional error is much larger for non-linear decay in tissues that pose a flux barrier to the morphogen at the boundary. In light of this new data, a physiological role of morphogen decay dynamics in patterning precision appears unlikely.

## Introduction

According to Wolpert’s famous French flag model [1], morphogen gradients encode readout positions *x*_*θ*_ via concentration thresholds *C*_*θ*_ = *C*(*x*_*θ*_), and differentiating cells base their fate decisions on whether the local morphogen concentration lies above or below such thresholds. Variations in the morphogen profile thus result in variations in the readout position. The accuracy of the spatial information carried by morphogen gradients can be quantified with the positional error, which is defined as the standard deviation of the readout positions over different gradient realizations [2]:

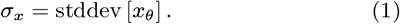

How the observed precision of tissue patterns arising from this principle is achieved, in spite of natural molecular noise in morphogen production, transport, decay, internalization, turnover and other sources of variability, is a key question in developmental biology [3, 2, 4].

Morphogen dynamics are often described by reaction-diffusion equations of the form [5]

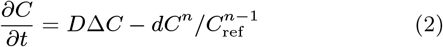

with morphogen concentration *C*, diffusion coefficient *D*, and decay rate *d. C*_ref_ is a constant reference concentration that we introduce to make all units independent of *n*. The exponent *n* models linear (*n* = 1) or non-linear (*n >* 1) decay of the morphogen. Non-linear decay would for instance ensue in case of cell lineage transport, when ligands interact with receptor clusters, or if ligand binding results in receptor upregulation, as is the case for several morphogens, most prominently for Hedgehog (Hh) [6–9]. Most reported morphogen gradient profiles have been fitted assuming linear decay (*n* = 1) [10–17]. For the FGF8 gradient in the developing mouse brain, *n* ≈ 4 has been reported [18].

Differences in sensitivity to a changing morphogen influx from the source into the patterned tissue have been argued to make gradients more robust if morphogen decay is non-linear, because they shift less when the morphogen influx is altered [6]. However, the gradients that result from non-linear decay also possess significantly shallower tails, relative to the higher concentration. Their usefulness for patterning has therefore been questioned [5], and it has remained unclear whether nonlinearity in the decay would in fact help achieving higher positional accuracy. Indeed, to first order, the positional error of variable gradients is inversely proportional to the magnitude of their slope [10, 2] according to

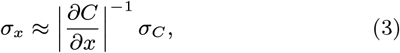

where *σ*_*C*_ is the standard deviation of local morphogen concentration and *x* denotes the patterning axis. This suggests that for patterning precision, the benefit of a smaller positional shift of gradients with *n >* 1 might be offset or even overcompensated by their flatter shape further from the source.

To investigate the physiological relevance of the mode of morphogen degradation, we employ here a recently developed numerical framework that allows us to analyse how physiological molecular noise translates into gradient variability, and thus, patterning precision [2, 19]. For reported molecular noise levels, the simulated gradients have previously been found to be sufficiently precise to yield the observed precision of developmental boundaries in case of linear decay [2]. We now extend the framework to analyse the impact of non-linear decay on gradient precision and find that it leads to a marginally lower positional error close to the source compared to linear decay. With only a small fraction of a cell diameter, the improvement is small and therefore hardly physiologically relevant. The effect of zero-flux conditions at the tissue boundary on non-linear decay, on the other hand, causes a drastically increased positional error in the patterning domain in comparison to linear decay.

## Results & Discussion

Before studying the effect of physiological variability, we start off with some theoretical considerations about the consequences of nonlinear decay for noise-free morphogen gradients.

### Qualitative difference between linear and non-linear morphogen decay

On one-dimensional, infinite patterning domains (*C*(*x*) → 0 as *x* → ∞), linear morphogen decay (*n* = 1) results in exponential steady-state gradient profiles (Fig. 1A) of the form [20]

**Figure 1:**
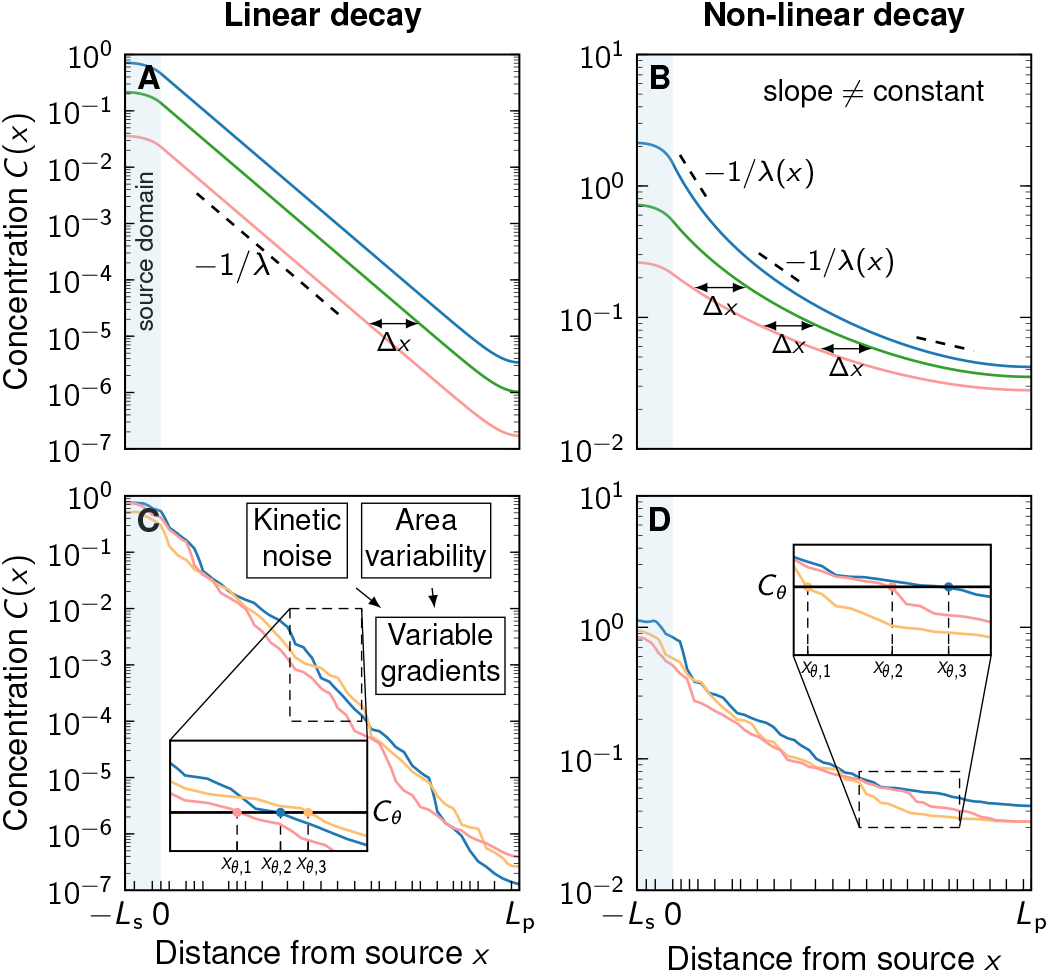
Comparison of linear and non-linear morphogen gradients. **A** Linear decay leads to exponential gradients. Changes in the gradient amplitude *C*_0_ (different colours) lead to a shift Δ*x* that is independent of the amplitude. **B** Non-linear decay (*n* = 2) leads to power-law gradients. The shift Δ*x* due to a change of *C*_0_ is amplitude-dependent. **A,B** With zero-flux conditions at the distal boundary, the shift is uniform in the patterning domain only away from that boundary. Cell boundaries are denoted by black ticks along the patterning axis. **C,D** Molecular kinetic noise and cell area variability leads to noisy gradients. For a fixed readout threshold *C*_*θ*_, variable gradients result in different readout positions *x*_*θ,i*_ (inset plots). Non-linear decay (**D**) leads to shallower gradients further in the patterning domain compared to linear decay (**C**).

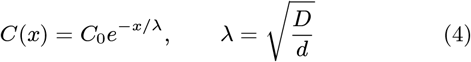

with *C*_0_ either the Dirichlet boundary condition at the source, or *C*_0_ = *j*_0_*λ/D* in case of a flux boundary condition at the source (− *D∂C/∂x*|_*x*=0_ = *j*_0_). With linear decay, influx and amplitude are thus proportional.

Non-linear decay (*n >* 1), on the other hand, results in shifted power-law gradients [6] (Fig. 1B) that can be written as

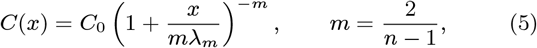

where

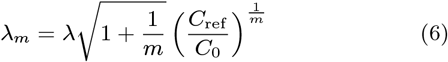

is a length scale determining the shift in the power law, and *C*_0_ = *C*(0) is the amplitude analogous to Eq. 4. As the linear decay is approached (*n* → 1), *m* diverges (*m* → ∞), the powerlaw length scale approaches the exponential length scale (*λ*_*m*_ → *λ*), and Eq. 5 converges to Eq. 4. For a flux boundary condition at the source, the morphogen amplitude is

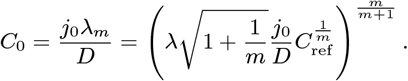

Amplitude and influx at the source boundary are thus not proportional for non-linear morphogen decay, unlike in the linear case. If one were to locally fit an exponential to the power-law gradient [6], *x*^−*m*^ ∼ exp [− *x/λ*(*x*)], one would observe the “gradient length” *λ*(*x*) to increase with the distance from the source according to *λ*(*x*) = *x/*(*m* ln *x*) (Fig. 1B).

The readout position at concentration threshold *C*_*θ*_ follows for linear decay from Eq. 4 as

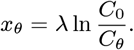

and for non-linear decay from Eq. 5 as

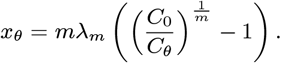

Real morphogen gradients are not deterministic but variable, and hence readout positions *x*_*θ,i*_ vary between different gradient realisations *i* for both linear and non-linear morphogen decay (Fig. 1C,D). Before we turn to the positional error this entails, we consider first the effect of a change on morphogen production levels on idealised, deterministic gradients. In response to a change in the morphogen amplitude from *C*_0_ to 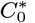, the readout position shifts along the patterning axis. For linear decay, this shift Δ*x* is independent of the absolute gradient amplitude *C*_0_ and depends only on the relative amplitude change, 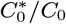, and the characteristic gradient length *λ*:

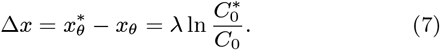

For non-linear decay, the shift is given by

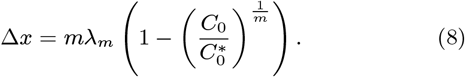

Since 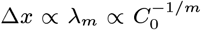, the shift increases with decreasing amplitude (Fig. 2A,B). This qualitatively distinguishes linear from non-linear decay. Alternatively, the shift may be expressed as a function of a change in morphogen influx from the source from *j*_0_ to 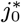. For linear decay, it simply reads

**Figure 2:**
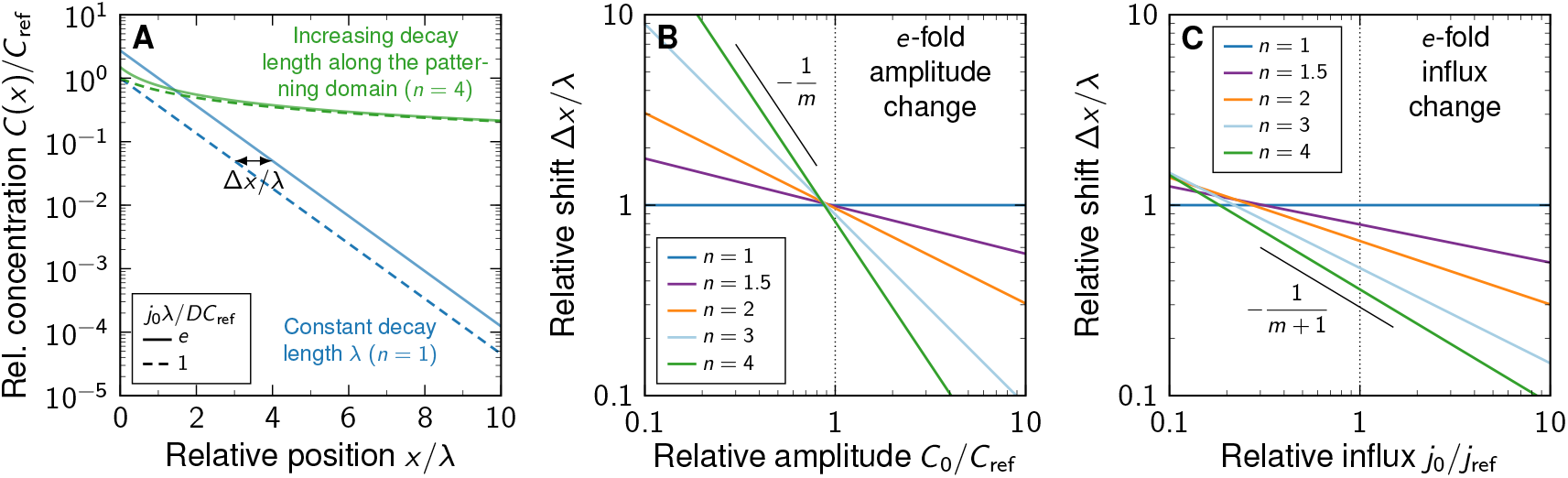
**A** Comparison of noise-free gradients arising from linear (blue) and non-linear (green) decay. A fold-change in the influx *j*_0_ from the source shifts the gradients by Δ*x*. **B** Positional shift of the morphogen gradient as a function of the amplitude and degree of non-linearity, for a fold-change in the amplitude, 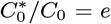. **C** Positional shift as a function of the influx and degree of non-linearity, for a fold-change in the influx, 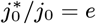.

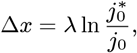

because flux and amplitude are proportional, making the shift again independent of morphogen levels. For non-linear decay, however, amplitude and influx are related as

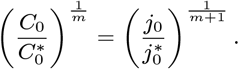

The resulting readout shift is therefore

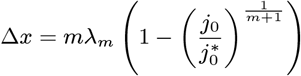

with a power-law length scale *λ*_*m*_ that can be expressed in terms of the influx *j*_0_ as

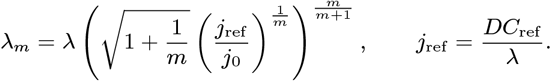

Therefore, since 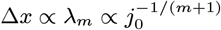, the shift also increases with decreasing influx (Fig. 2A,C), albeit slower than with the amplitude.

It has previously been argued [6] that the circumstance that the power-law shift vanishes at sufficiently large influx values (Δ*x* → 0 as *j*_0_ → ∞for *n >* 1) would lead to more robust patterning, because the readout position becomes independent of the influx in this limit when the morphogen decay is non-linear. In this discussion, molecular noise was included only in the form of a fold-change in the amplitude or influx. We now extend this deterministic view by taking molecular variability of the gradients into account, and demonstrate numerically that the positional error of noisy morphogen gradients does not significantly improve with non-linear decay. In the contrary, if the morphogen cannot leave the patterning domain opposite of the source, the power-law gradients become shallow in a substantial part of the domain, leading to reduced positional accuracy with non-linear decay.

### Noisy gradient model

To study the impact of decay non-linearity on the precision of noisy morphogen gradients, we simulated steady-state diffusion on a one-dimensional cellular domain composed of a source of length *L*_s_ and a patterning region of length *L*_p_ (Fig. 1C,D). Eq. 2 was extended by a morphogen production term in the source, resulting in

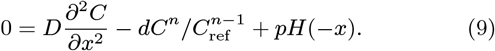

Here, *H*(*x*) is the Heaviside function, ensuring that production at rate *p* only occurs in the source (*x <* 0). No-flux boundary conditions were used at both outer ends of the domain:

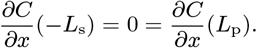

We generated large numbers of variable morphogen gradients by numerically solving Eq. 9 with kinetic parameters *p, d* and *D*, and cell areas *A* independently drawn from log-normal distributions for each cell in the domain, as described before [2, 19]. The diffusion coefficient, *D*, and the degradation rate, *d*, set the steady-state patterning length scale, 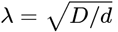, and we report positional quantities relative to the average *λ* or the average cell diameter, which in turn is chosen to be a fixed multiple of the average *λ*. Thus, our results are independent of the absolute values chosen for *D* and *d*.

We express the mean value and standard deviation of a parameter *q* by *µ*_*q*_ and *σ*_*q*_, respectively. Based on measurements of the Hedgehog morphogen gradient in the *Drosophila* wing disc and mouse neural tube [12, 15, 2], we used *µ*_*D*_ = 0.033 µm^2^/s, *µ*_*λ*_ = 20 µm. Other specific values would not change the results reported here, which are for the steady state, but would only alter the timescale it takes for the steady state to be reached. We furthermore set 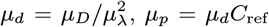, *µ*_*p*_ = *µ*_*d*_*C*_ref_, as average ki-netic parameters, where *C*_ref_ = 1 arb. units to normalise the concentrations.

The noise-to-signal ratio in each quantity *q* is given by the corresponding coefficient of variation, CV_*q*_ = *σ*_*q*_ */µ*_*q*_. Reported physiological noise levels in morphogen production, decay, and transport differ between morphogens and tissues, but are around CV_*p,d,D*_ ≈0.3 [2], which we use to define the distribution widths of the kinetic parameters.

The widths and cross-sectional areas of cells vary in all layers along the apical-basal axis [21]. Most quantifications have been carried out on the apical surface. One of the highest reported values for the apical area CV is found in the vertebrate neural tube (CV_*A*_ ≈0.9) [22–24], but most values are considerably lower [25]. We therefore used CV_*A*_ = 0.5 in all simulations unless specified otherwise. Random cell areas were drawn from a log-normal distribution with the specified CV_*A*_ and mean cell area [19]

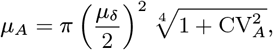

allowing us to control the mean cell diameter *µ*_*δ*_. The individual random cell areas *A* were then transformed to cell diameters *δ* according to 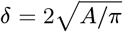, and the spatial axis was discretized into cellular sub-intervals accordingly. We used a patterning domain length of 200 cells (*L*_p_ = 200*µ*) and a source domain length of 5 cells (*L*_s_ = 5*µ*_*δ*_), unless otherwise stated.

Whether cells average the morphogen signal over their entire cell surface, beyond their cell surface via a cilium, or read out the signal at a single point, has little effect on the readout precision [19]. We therefore only analysed the case where cells average the morphogen signal over their cell surface (their diameter in the 1D model here).

### Impact of non-linear decay on gradient precision

We previously showed that in case of linear decay, there is a negligible impact of cell area variability as long as CV_*A*_ *<* 1 [19]. We now find that this holds similarly for non-linear decay (Fig. 3A), justifying the use of a fixed CV_*A*_ = 0.5 in the remainder of our analysis.

**Figure 3:**
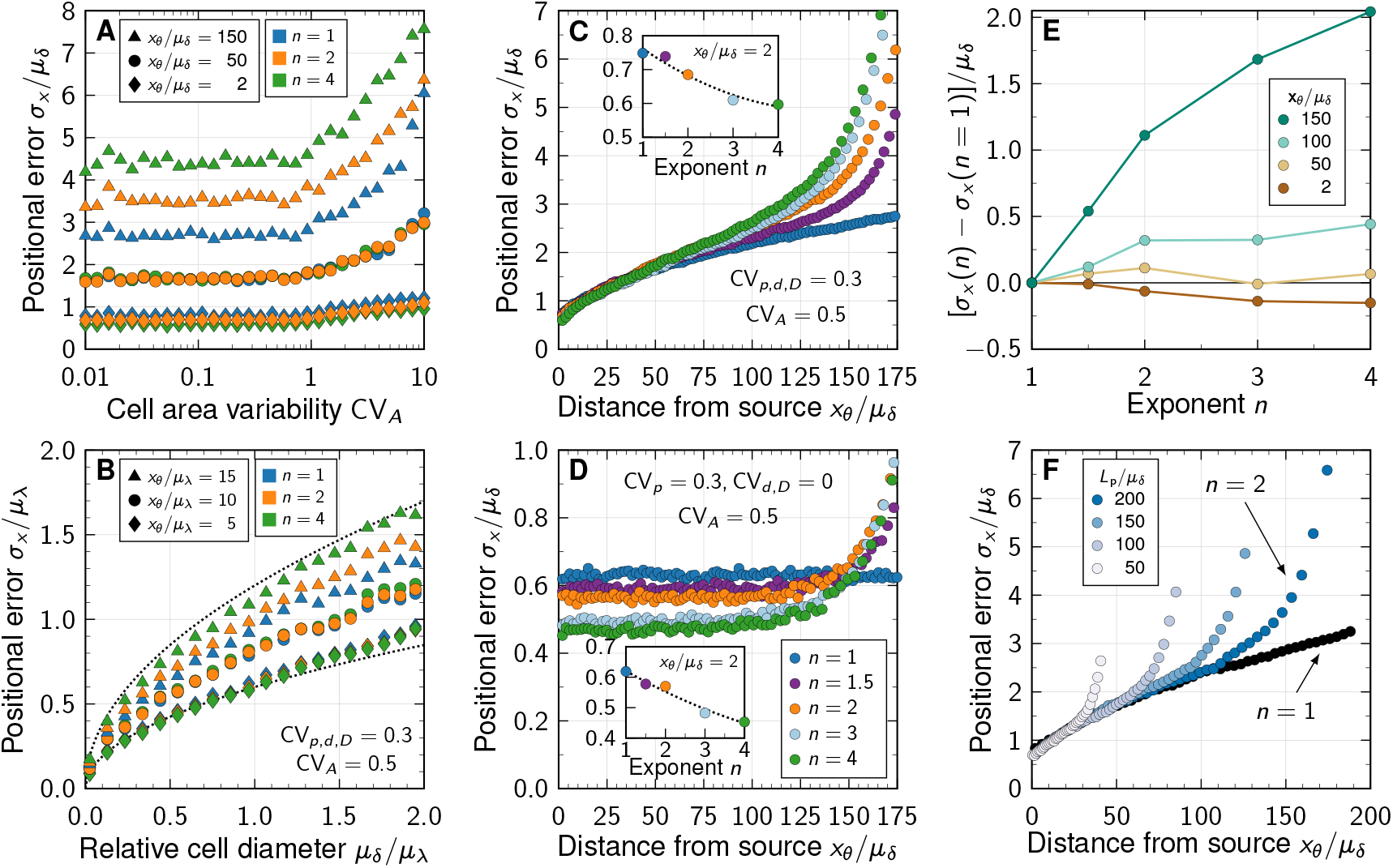
Impact of non-linear decay on gradient precision. **A** Physiological variability in the cross-sectional cell areas has no significant impact on gradient precision. The positional error *σ*_*x*_ is plotted in units of the mean cell diameter *µ*_*δ*_ at different readout positions in the patterning domain (symbols), and for different degrees of non-linearity (colours). **B** The positional error increases with the square root of the cell diameter, irrespective of *n*. Dotted lines show 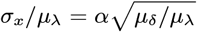 for *α* = 0.6, 1.2 for reference. *L*_p_ = 100*µ*_*δ*_. **C** Non-linear decay leads to a marginally lower positional error close to the morphogen source. Inset plot shows *σ*_*x*_*/µ*_*δ*_ at a distance of two cells from the source as a function of decay non-linearity. With a no-flux boundary at *x* = *L*_p_, the shallowness of gradients from non-linear decay lets the positional error increase strongly far from the source. **D** Variability in the production rate alone has no effect on the positional error along the domain for linear decay (blue). The stronger the non-linearity, the smaller the positional error close to the source (inset). Far from the source, the positional error increases rapidly with non-linear decay. **E** Difference between the positional error for *n >* 1 and for *n* = 1 relative to the mean cell diameter, at fixed readout positions (colours). **F** Effect of finite patterning domain size. The positional error increases close to the distant zero-flux boundary in case of non-linear decay (shades of blue, *n* = 2). Patterning remains precise across a larger distance in the case of linear decay (black, *n* = 1). In all panels, each data point represents the mean from 10^3^ independent simulations. Standard errors are smaller than the symbols.

Much as for linear decay [19], the positional error scales with the square root of the mean cell diameter also for non-linear decay (Fig. 3B). Small cell diameters, as observed in all known tissues that employ gradient-based patterning [19], are therefore important for high spatial precision also in case of non-linear decay. For the remainder of this study, we fix the average cell diameter at a fourth of the exponential gradient length, *µ*_*δ*_*/µ*_*λ*_ = 1*/*4, as found in the developing mouse neural tube [26, 15].

The positional error increases from less than one cell diameter close to the source to about two cell diameters at a distance of 75 cell diameters away from the source (Fig. 3C). Close to the distant domain boundary opposite of the source, where a no-flux condition was imposed, the positional error rapidly increases for non-linear decay, while remaining relatively low for linear decay. If only the production in the source is varied (CV_*p*_ = 0.3, CV_*d,D*_ = 0), the positional error remains constant as the readout distance from the source increases, but increases again sharply close to the distant end in case of non-linear decay (Fig. 3D). But even for strong non-linearity (*n* = 4), the positional error remains in the sub-cellular range when only production noise is considered, as long as the readout position is further than about *λ* away from the distal end.

Independent of whether all parameters are varied or only the production rate, the positional error drops in close vicinity to the source with stronger non-linearity in the decay (insets of Fig. 3C,D). However, with less than 20% of a single cell diameter from *n* = 1 to *n* = 4, the effect is likely too small to be physiologically relevant. Further away from the source, linear decay yields a smaller positional error than non-linear decay (Fig. 3D–E). No matter how long the patterning domain is, non-linearity always increases the positional error as the distal tissue boundary is approached (Fig. 3F).

What then causes the increased positional errors with non-linear decay near the distal domain boundary? A zero-flux boundary condition there implies shallower gradients than on infinite domains: *C*^*′*^(*x*) → 0 as *x* → *L*_p_. This effect occurs irrespective of *n*, but the spatial range over which the gradient flattens (and thus deviates from the pure exponential and shifted power-law forms for infinite domains, Eqs. 4 and 5) increases with *n*. By virtue of Eq. 3, non-linear decay thus leads to greater positional errors at readout positions in the vicinity of the distal boundary compared to linear decay.

In summary, our computer simulations of noisy morphogen gradients suggest that it is insufficient to quantify gradient robustness and patterning precision by considering variability in the morphogen production alone. Moreover, if the morphogen cannot exit the patterning domain opposite of the source, shifted power-law gradients that result from non-linear morphogen decay flatten over a significantly larger range than exponential gradients, leading to increased positional errors. The gain in positional accuracy close to the source for non-linear decay is negligible and therefore barely physiologically relevant. Overall, exponential gradients lead to more robust patterning.

### Impact of boundary condition at the source

Given the impact of the distal domain boundary, we wondered whether the representation of the morphogen source by either a spatial production domain (Fig. 4A), by a flux boundary condition −*DC*^*′*^(0) = *j*_0_ (Fig. 4B) as used by Eldar et al. [6], or by a fixed gradient amplitude *C*(0) = *C*_0_ (Fig. 4C) would affect the positional error predicted by the model. While there are small quantitative differences, the gradient shapes (Fig. 4A–C) and positional errors (Fig. 4D–F) are overall very similar.

**Figure 4:**
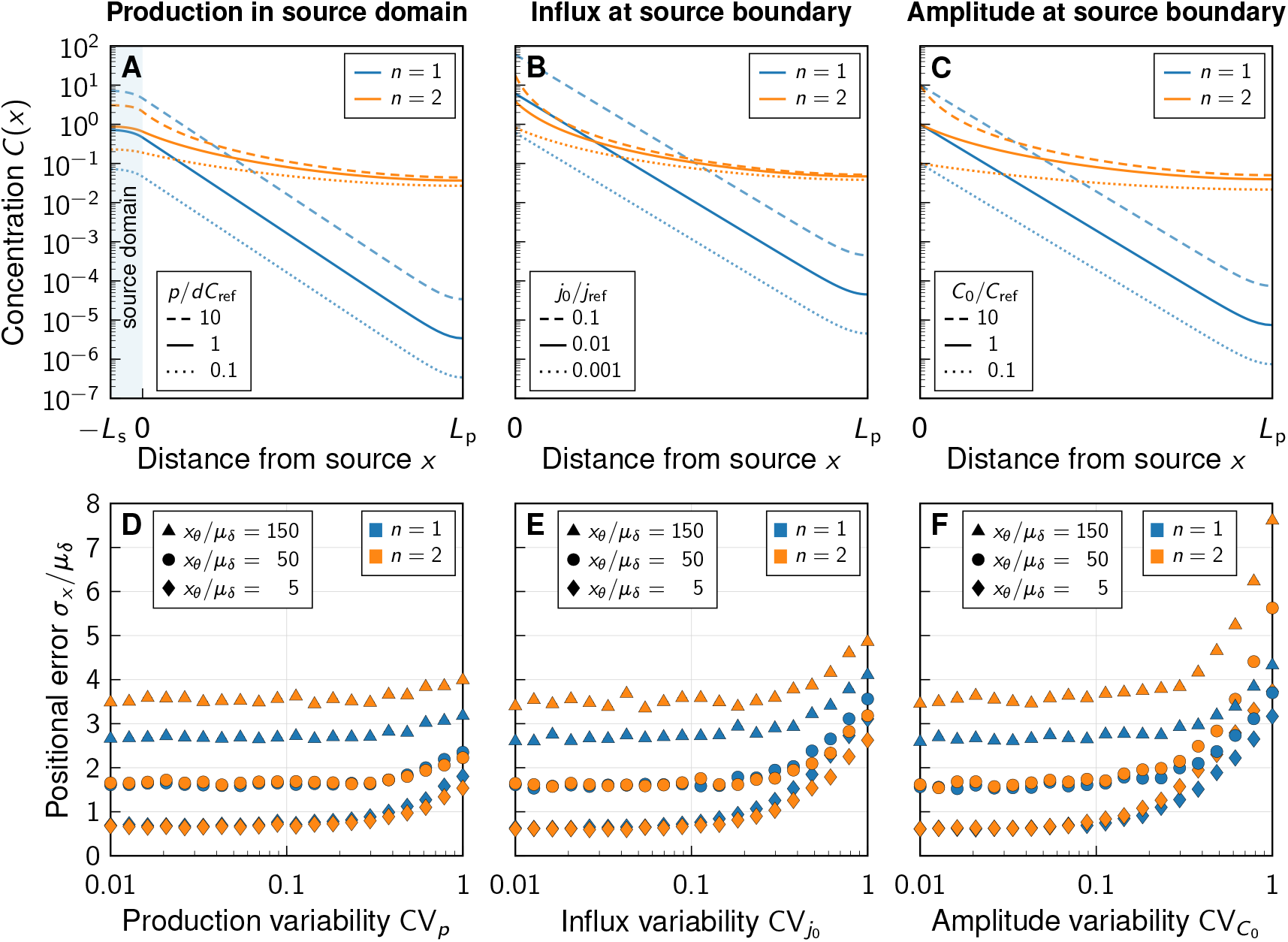
Impact of the boundary condition (BC) at the source. **A–C** Noise-free gradient shapes when the morphogen is either secreted in a source domain at rate *p* (Eq. 9) (A), with flux BC, − *D∂C/∂x* _*x*=0_ = *j*_0_(B), or Dirichlet BC, *C*(0) = *C*_0_ (C). No-flux BC were imposed at at the far end of the tissue (at *x* = *L*_p_). **D–F** Positional error as a function of morphogen abundance variability, at different readout positions (symbols) and degrees of non-linearity (colours). Greater variability in the morphogen production rate (D), influx (E), and gradient amplitude (F) leads to a larger positional error above a certain threshold variability CV ⪆ 0.1–0.3. Kinetic variability was fixed at CV_*p,d*_ = 0.3 (except for CV_*p*_ in D). Further parameters: 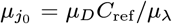 (E), 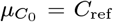 (F). In panels D–F, each data point represents the mean from 10^3^ independent simulations. Standard errors are smaller than the symbols.

As we increase the variability in the production rate via CV_*p*_ (Fig. 4D), in the influx from the source via 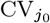 (Fig. 4E), or in the gradient amplitude at the source boundary via 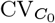 (Fig. 4F), we find the smallest increase in the positional error for the production rate and the largest increase for the gradient amplitude. Neumann or Dirichlet boundary conditions thus overestimate the positional error when the variability in the source is high. Instead of using such boundary conditions, a spatial source domain should explicitly be modeled, where applicable. With the physiological values CV_*p*_ ≈ 0.3 and 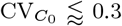 [2], however, variability in the morphogen production plays merely a subordinate to moderate role in the overall gradient variability. Molecular noise in morphogen degradation and diffusivity dominates the patterning precision.

### Impact of the morphogen source strength

As the gradient amplitude determines the sensitivity of the read-out position to amplitude changes for non-linear decay (Eqs. 6 and 8) but not for linear decay (Eq. 7), the average morphogen production rate is expected to affect the patterning accuracy in the case of non-linear decay, but not for linear decay. We put this theoretical prediction to the test by varying the mean relative production rate, the mean influx from the source, and the mean morphogen amplitude in the three different simulated morphogen production models. Changes in these parameters have no effect on the positional error if morphogen degradation is linear, which is consistent with the theory (Fig. 5A–F, blue lines). With non-linear decay, on the other hand, we indeed observe the positional error to be highly dependent on morphogen abundance. This dependency, however, is not well approximated by simple power laws 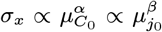, such as for the shift 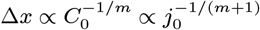 discussed in the context of noise-free gradients above. Precision arguments previously brought forward for deterministic morphogen gradients [6] do not appear to directly quantitatively translate to the positional error in settings where cell-to-cell variability is included, and where morphogen production remains at physiological levels.

**Figure 5:**
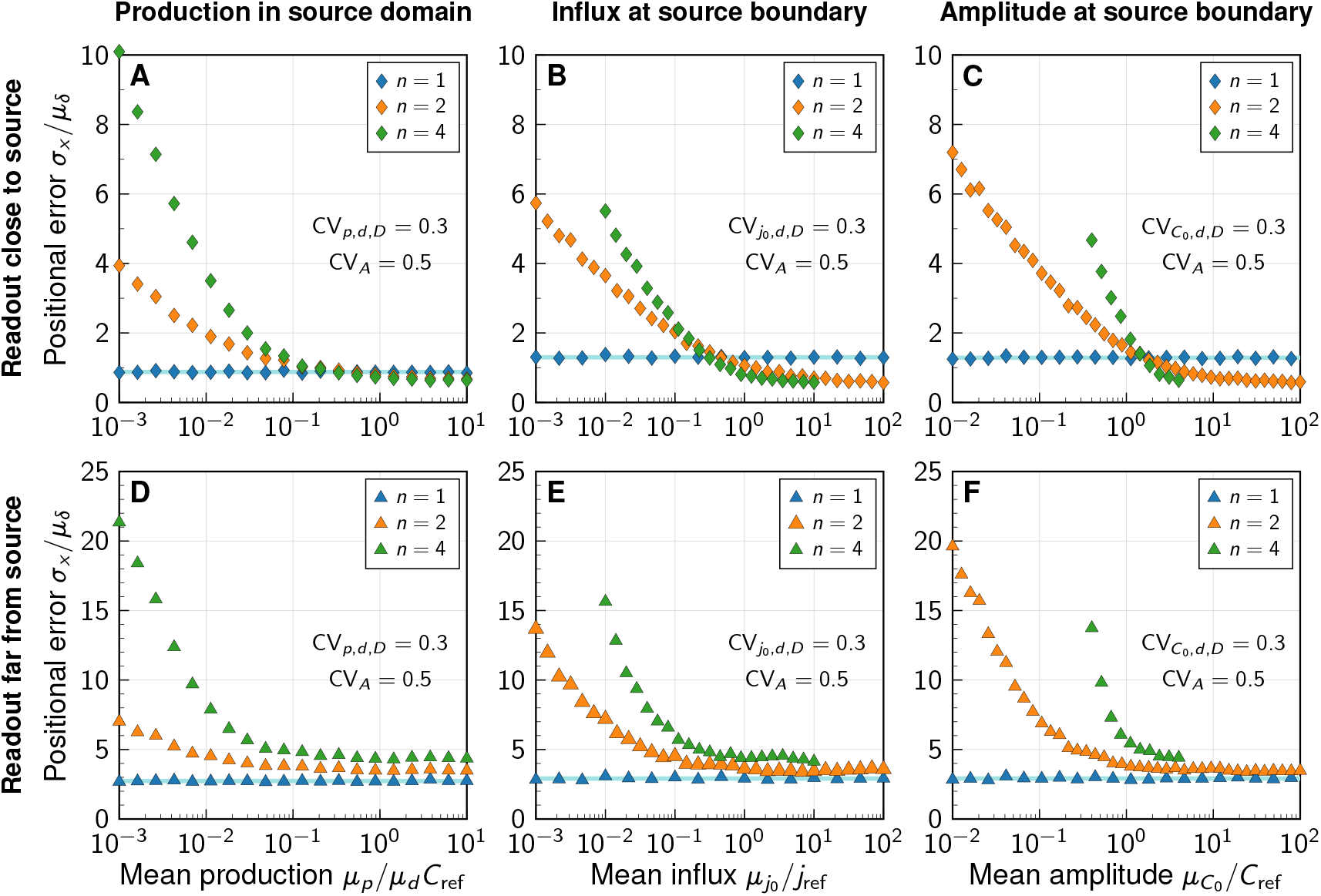
Impact of the morphogen source strength. Numerically obtained spatial patterning accuracy in units of average cell diameters *µ*_*δ*_ at different positions in the tissue (symbols) and for different degrees of non-linearity (colours). **A–C** Readout close to the source, at *x*_*θ*_ = 5*µ*_*δ*_; **D–F** Readout far from the source, at *x*_*θ*_ = 150*µ*_*δ*_. Morphogen production scenarios are identical to Fig. 4: Production in a source domain with morphogen-secreting cells (A,D), with a morphogen influx from the source at the source boundary (B,E), and with a specified morphogen concentration at the source boundary (C,F). Very low (high) influxes or amplitudes lead to flat (steep) gradients at strong decay non-linearity, limiting the parameter range over which the positional error can be reliably determined for *n* = 4 (B,C,E,F). In all panels, each data point represents the mean from 10^3^ independent simulations, with standard errors smaller than the symbols.

For high morphogen supply levels, non-linear decay leads to a smaller positional error close to the source (Fig. 5A–C). The effect is, however, substantially less pronounced in the model that includes a spatial morphogen source domain (Fig. 5A) than in those that do not (Fig. 5B,C), highlighting once again the limitations of the latter. With an explicit source domain, non-linear decay yields only marginally more spatial accuracy, when production is high (*p/dC*_ref_ ⪆ 0.4). Lower production levels increase the positional error close to the source substantially in all three models, reaching several cell diameters, for *n >* 1. The gradients effectively flatten out at low production, reducing their usefulness for spatial tissue patterning. The stronger the non-linearity in the degradation, the more pronounced this loss of patterning precision.

Further away from the source, the benefit of non-linear decay is lost entirely, and exponential gradients remain more precise than shifted power-law gradients also at high morphogen supply levels (Fig. 5D–F).

In summary, simplified models without explicit representation of morphogen-secreting cells overestimate the beneficial impact of non-linear decay on patterning precision. In all models considered here, the benefit of non-linear morphogen decay is restricted to a close vicinity of the morphogen source, where patterning precision is high anyway [2] and may thus not be as critical for robust development, and to a regime of very strong morphogen production. Further into the tissue, and at moderate morphogen abundance, linear decay yields more accurate patterning.

## Conclusion

Non-linear morphogen decay was proposed as a potential precision-enhancing mechanism for tissue patterning in the seminal theoretical work by Eldar et al. [6] in a deterministic setting, where morphogen gradients are devoid of noise. Here we have explored this idea with a stochastic model, taking noisy gradients into account, as they arise from cell-to-cell variability in morphogen kinetics. The surprising outcome of our quantitative analysis is that, while a small advantageous effect indeed exists near the morphogen source, this gain is outweighed by a substantial loss of precision in the spatial information that signalling gradients provide to cells in the interior and distal parts of a patterned tissue when morphogen decay is non-linear. In tissues that pose a diffusion barrier to the signalling molecule at their boundary, shifted power-law gradients that emerge with self-enhanced degradation, flatten out over a substantial portion of the spatial domain, whereas exponential gradients remain more graded (Fig. 1). This leads to greater spatial precision with linear decay (Fig. 3), and is contrary to the original expectation [6].

This long-range boundary effect is not the only reason why linear morphogen decay is favourable for precise pattern formation. The positional error, which is the decisive quantity that measures the spatial accuracy with which cells can determine their location in the pattern, and ultimately their fate in differentiation, is highly sensitive to morphogen supply levels when morphogen decay is non-linear, but largely insensitive when decay is linear (Fig. 5). This implies that patterning is more robust to variations in the size and strength of the morphogen-secreting source, if decay is linear. These results challenge the established view that power-law gradients buffer fluctuations in morphogen production [6]. We find that the positional error behaves in the opposite way, buffering production fluctuations only with linear, but not with non-linear decay. From an evolutionary perspective, the linear case may be favoured, as patterning precision is unaffected by changes in the size and kinetics of the morphogen-secreting source only if *n* = 1.

Our study demonstrates that a stochastic approach is required to quantify patterning precision of real noisy gradients. Moreover, we find the positional error to be overestimated in simplified models that replace the morphogen-secreting cells by a Neumann or Dirichlet boundary condition (Fig. 4). Based on this, we recommend to include an explicit representation of the source in future theoretical or numerical work on the subject, as we did with Eq. 9.

Distinguishing exponential gradients from shifted power laws can be very difficult in practice, as they can appear similar over the short distances over which they can be reliably measured with classical imaging techniques. The FGF8 gradient in the developing mouse brain is the only case we are aware of where *n >* 1 has been reported robustly [18], and whether this is linked to patterning precision in any way remains unclear. Available gradient data in other systems, such as Sonic Hedgehog and Bone Morphogenetic Protein in the neural tube [16], is too variable to confidently reject the hypothesis that *n* = 1. Most further reports of morphogen gradient shapes [10–17] are consistent with exponentials within measurement accuracy. New measurement techniques are needed to determine whether non-linear decay is at work in the formation of known morphogen gradients during development. In light of our findings, a physiological role of non-linear ligand decay in patterning precision appears implausible. If anything, our data suggest an overall advantage of linear decay, also considering the evolutionary aspect of tissue size and protein synthesis rate differences between species.

## Code Availability

The source code is released under the 3-clause BSD license. It is available as a public git repository at https://git.bsse.ethz.ch/iber/Publications/2022_adelmann_vetter_nonlinear_decay.

## Acknowledgements

This work was partially funded by SNF Sinergia grant CR-SII5 170930.

## Competing Interests

None declared.

## Author Contributions

RV & DI conceived the study, RV derived the theory, JA and RV developed the numerical framework, JA carried out the simulations, JA and RV produced the figures. All authors wrote the manuscript.

## Notes

https://git.bsse.ethz.ch/iber/Publications/2022_adelmann_vetter_nonlinear_decay

### Competing Interest Statement

The authors have declared no competing interest.

